# DBD-RCO: Derivative Based Detection and Reverse Combinatorial Optimization to improve heart beat detection for wearable devices

**DOI:** 10.1101/118943

**Authors:** Andrea Bizzego, Cesare Furlanello

## Abstract

The diffusion of wearable sensors enables the monitoring of heart physiology in real-life contexts. Wearable technology is characterized by important advantages but also by technical limitations that affect the quality of the collected signals in terms of movement artifacts and presence of noise. Therefore specific signal processing algorithms are required to cope with the lower quality and different characteristic of signals collected with wearable sensor units. Here we propose and validate a pipeline to detect heartbeats in cardiac signals, extract the Inter Beat Intervals (IBI) and compute the Heart Rate Variability (HRV) indicators from wearable devices. In particular, we describe the novel Derivative-Based Detection (DBD) algorithm to estimate the beat position in Blood Volume Pulse (BVP) signals and the Reverse Combinatorial Optimization (RCO) algorithm to identify and correct IBI extraction errors. The pipeline is first validated on data from clinical-grade sensors, then on a benchmark dataset including examples of movement artifacts in a real-life context. The accuracy of the DBD algorithm is assessed in terms of precision and recall of the detection; error in the IBI values is quantified by root mean square error. The reliability of HRV indicators is evaluated by the Bland-Altman ratio. The DBD algorithm performs better than a state-of-art algorithm for both medical-grade and wearable devices. However, as already found in similar studies, worse reliability is found with the BVP signal in computing frequency domain HRV indicators, in particular with wearable devices.

## 1 Introduction

The diffusion of wearable sensors enables the monitoring of heart physiology in real-life contexts. Besides several advantages wearable technology is characterized by technical limitations that affect the quality of the collected signals in terms of movement artifacts and presence of noise. Therefore the signal processing procedures require specific algorithms able to cope with the lower quality and different application domain of the signals collected by wearables. In this paper we introduce and validate a pipeline for Heart Rate Variability (HRV) analysis on signals acquired with Wearable Devices (WDs). Currently two main sensing technologies are available to acquire the cardiac activity with WDs: electrocardiography or photoplethysmography (PPG) sensors. The former is used to acquire the electrocardiogram (ECG), while the PPG sensors detect the Blood Volume Pulse (BVP).

A typical HRV analysis pipeline (see Figure 1) is composed of three stages [10]: (a) preprocessing; (b) beat detection and (c) computation of indicators, Typically applied on ECG signals. However, BVP signals can also be used [15]: from the signal processing point of view the main difference is in the second stage, where, due to the different waveform of BVP compared to ECG, specific algorithms have been defined to correctly estimate the beat position.

**Figure 1:**
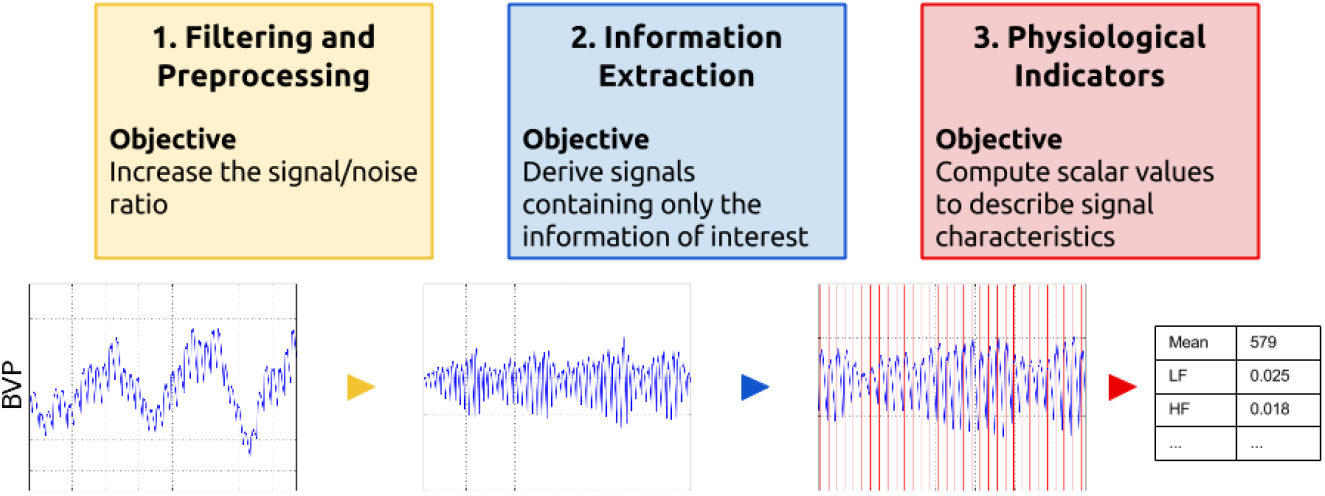
Three steps of physiological signal processing (top) and an example on a PPG signal (bottom): original signal (left), after preprocessing (middle left), result of beat detection (middle right) and computed physiological indicators of Heart Rate Variability (right).

### Preprocessing

The preprocessing part aims at increasing the signal-noise ratio (SNR) by filtering, detrending and removal of undesired components. In particular, several algorithms and technical solutions have been adopted to deal with moving artifacts (see, for instance [5, 13, 14, 17, 19] for a review), but complete removal is still an open issue due to the impulsive and time-dependency characteristic of body movements.

As an alternative, it has been proposed to estimate the amount of noise and reject those portions where this is too high [17, 11, 9]. However, in this paper we do not discuss denoising, focusing instead on the second stage of the pipeline: detection of heart beats and then computation of Inter Beat Intervals (IBIs).

### Beat detection

The detection of the beat position in an ECG signal corresponds to the identification of the R peak, which is usually well recognizable due to its higher amplitude and characteristic impulsive shape. Peaks can be identified using different algorithms (see for instance [8]) also for real-time processing (see [3]); however identification accuracy is weakened by high frequency noise and trends that should have been removed in the preprocessing stage.

A BVP signal does not provide a wave portion corresponding to the R peak of the ECG signal; in addition, beat detection is made difficult also by the variability due to posture or even individual physiological characteristics [15]. Indeed, a different family of algorithms is used to identify the correct beat position from PPG [1, 16, 12]. In general, it emerges that a reliable and general method is still missing.

Mostly, such algorithms are usually proposed for use in diagnostics and health care; in the latter case, the interest is a continuous HR monitoring and thus a rougher measure is adopted compared to HRV analysis (heart rate in beats per minutes vs measuring distance between beats at higher resolution of milliseconds). Further, only few algorithms have been proposed appositely for analysis of signals acquired with WDs. They present usually even higher variability in terms of waveform shape and thus algorithms need to be robust to time-variant characteristics of the signal and variations of amplitude with these sensors.

### Computation of HRV indicators

The first step in order to compute HRV indicators, is detecting heart beats from the cardiac signal, then derive the distance between consecutive beats (IBI). HRV indicators are used to describe a particular aspect of the heart physiology or activity of the autonomic system (for a complete review see [10]).

Here we propose a signal processing pipeline suited for signals acquired with WDs, which could serve as a reference methodology for future papers and comparison of new algorithms (Section 2). In particular, we describe the Derivative Based Detection (DBD) algorithm for beat detection on BVP signals and the Reverse Combinatorial Optimization (RCO) for correction of misdetection errors. We compare the DBD with the automatic beat detection algorithm [1] showing that DBD is more robust and appropriate for signals acquired with WDs and correction by RCO further improves the results. Section 3 describes the two datasets we used to test the pipeline: a) the Fantasia dataset [7] and b) our WCS dataset, designed to compare signals from clinical (Thought Technology FlexComp Infinity) and wearable devices (Empatica E4 and ComfTech HeartBand). We present and discuss the result in terms of beat detection errors and differences in the computed HRV indicators (Section 4 and 5).

## 2 Beat detection pipeline

In this section we present the pipeline for beat detection on cardiac signals to extract the IBI signal. The pipeline addresses the issues originated by use of acquired wearable technologies, however it can be applied on signals from medical-grade devices, where it is expected to give even better performances (see Section 5).

In particular, we present the DBD algorithm for estimation of beat position in BVP signals and the RCO algorithm for errors detection and correction. The interest of the DBD algorithm is that most of WDs embed a PPG sensor to acquire cardiac signals. We omit to discuss the preprocessing step (for instance band-pass filtering to remove trends and high frequency noise) as already extensively discussed in literature.

### 2.1 Oversampling

Effects of lower sampling frequency on the HRV indicators can be mitigated by signal interpolation [4]. The first step in the pipeline is therefore an oversampling to decrease the error in the estimation of the beat position. We apply a cubic spline interpolation, with an output sampling frequency of at least 1000 Hz to estimate distances between beats with 1 ms resolution.

### 2.2 Energy estimation

Several methods have been proposed to deal with artifacts associated to body movements but in case of high dynamic movements or severe signal corruption (for instance due to electrodes disconnection or displacement) they might be ineffective [17]. Further, the presence of artifacts in the signal will always impact on the error of estimation of the beat position. For this reason, beside addressing the removal of artifacts, it is important to obtain a measure of reliability of the estimates. This becomes fundamental in online applications (e.g. when shortterm or real-time indicator triggers a decision logic), or when processing large amounts of streaming flows which would make practically unfeasible to manually check the results of beat position estimation.

To this aim we propose to use the Energy of the Signal Derivative:

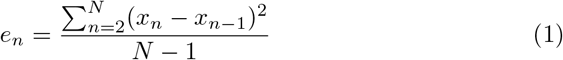

where *x_k_* is the value of *k*-th sample of the signal and *N* is the total number of samples.

To consider for local variations of the noise the energy (1) should be iteratively computed within a windowing process that runs along the whole signal:

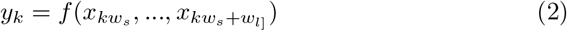

where *k* is the current windowing iteration *w_s_* is the number of samples the window shifts at each iteration and *w_l_* is the number of samples of the window.

In absence of artifacts, *e_n_* depends only on the dynamic of the signal which can be considered approximately time invariant. When artifacts corrupt the signal their contribution will increase the value of *e_n_* which can therefore be used to estimate the presence and magnitude of artifacts A threshold value should be defined to identify the noisy portion of signals that would provide an unreliable estimation of the beat position and, consequently, of the computed HRV indicator.

Note that, when the wearable devices embeds also an accelerometer, *e_n_* can be computed from the acceleration signal to provide a more reliable quantification of artifacts due to body movement.

### 2.3 The DBD algorithm

The DBD algorithm estimates the position of the percussion peak in signals acquired with an optical sensor and it is designed to be robust when applied on signals from WDs which present high variability of waveform shapes and amplitudes.

As for other algorithms proposed in literature (e.g. [1, 12]),the DBD algorithm adopts a sequence of processing steps. We use three main stages, with an additional stage for the automatic identification of detection errors (RCO). An overview of the DBD procedure is presented in Figure 2.

**Figure 2:**
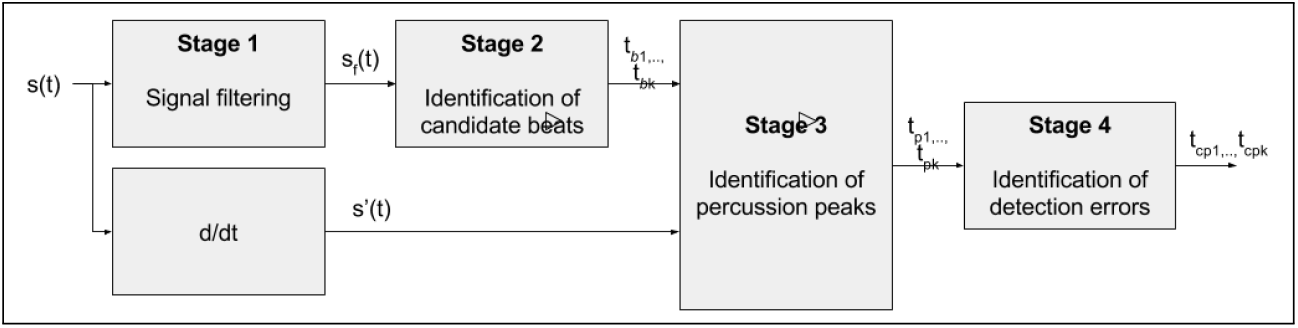
Overview of the DBD algorithm with the four processing stages.

The algorithm is regulated by the parameter *f*_max_ which is the expected maximal heartbeat frequency. We empirically set *f*_max_ to 2 Hz, corresponding to 120 beats per minute). However, the parameter can be changed accordingly to physiological condition (e.g. sleep) or age of individual (e.g. infant, newborn). In the following subsections a detailed description for each stage of the algorithm is given.

#### DBD Stage 1: Signal filtering

The first stage aims at filtering the original signal to extract the approximate beat positions. This information will be used in the following steps to target the detection of the percussion peak. We use a low pass infinite impulse response filter (*f_pass_* = 1.2*f*_max_, *f_stop_* = 3*f*_max_) to filter out high frequency components of the signal.

#### DBD Stage 2: Identification of candidate beat position

On the filtered signal from stage 1 we identify the candidate beats by adaptive peak detection. First the local range of the signal is estimated by windowing (window width: 1.5(*f*_samp_)/*f*_max_, window shift: *f_samp_*/*f*_max_) and evaluating the range for each window, then interpolating according to the original sampling frequency of the signal. The local range of the signal is used to modulate the peak detection: a local maximum 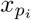 is considered valid if the difference with the following local minimum is greater than half of the range of the signal at the local minimum.

#### DBD Stage 3: Identification of percussion peak position

For each peak instant detected in stage 2 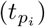 we extract the 250 ms before 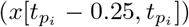 which is the candidate portion 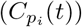 where to locate the percussion peak. We compute the derivative of the candidate portion 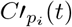 where we detect the maximum 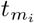 corresponding to the steepest instant of the rising part of the pulse and the instant 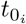 following 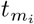, corresponding to the first local minimum of 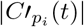.

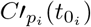 is expected to be equal to 0 and is considered the instant of the percussion peak. Figure 3 shows the steps to identify 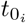 on a beat pulse. The IBI signal is then computed as difference between consecutive percussion instants.

**Figure 3:**
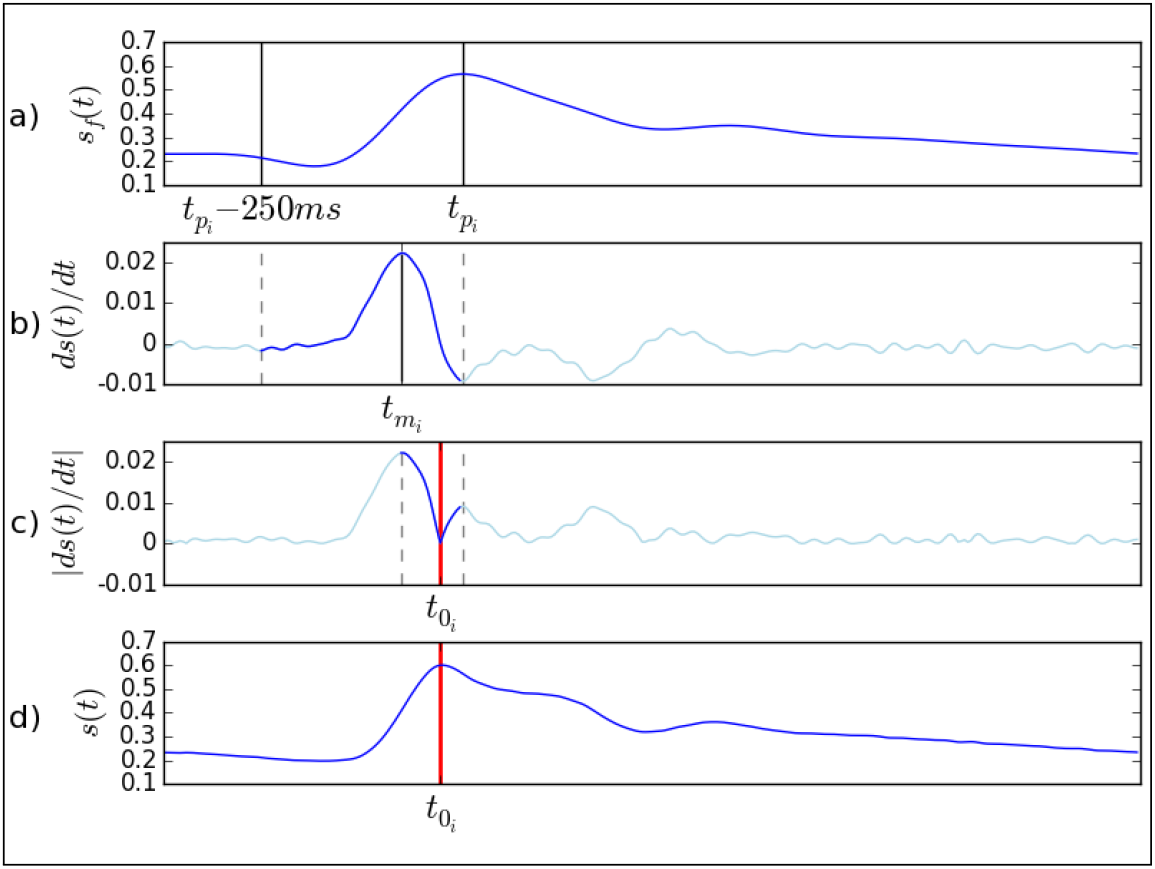
Steps of stage 3 to identify 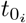. From top to bottom: a) identification of the maximum 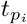 and 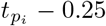 on the filtered signal *x_f_* (t) ; b) identification of the maximum of the derivative of the signal *x_n_* − *x*_*n*−1_ in the interval 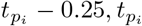; c) identification of the local minima of |*ds*(*t*)/*dt*| in the interval 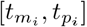, corresponding to 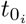, the percussion peak in the original signal (d).

### 2.4 The RCO algorithm

The RCO algorithm aims at identifying and correcting errors in the IBI signal in particular when the correct beat position is not detected because of a false peak before the true peak. It depends only on the IBI signal and thus it can be employed also to correct beats detected on ECG signals.

#### Adaptive Outlier Detection

The basic algorithm of the RCO is the Adaptive Outlier Detection (AOD). The AOD uses a fixed size cache vector *IBI_c_* = (*ibi*_1_*, ibi*_2_,*…, ibi_k_)* to store the last *k* valid IBI values, in order to adapt to IBI variability. The size *k* is empirically set to 5 and the cache is initialized with the median value of the IBI signal. Outliers detection is also regulated by the sensitivity parameter *ϕ*, which is empirically set to 0.25. However, smaller values of *ϕ* and greater values of *k* improve precision.

A detected beat is considered valid if its corresponding IBI value is within the interval [(1+*ϕ*)*ibi_medion_*,(1+*ϕ*)*ibi_medion_*], where *ibi_median_* is the median of the values in *IBI_c_.* When a new valid beat is detected, its IBI value is used to update *IBI_c_* using the First-In-First-Out rule. *IBI_c_* is re-initialized when *k* consecutive non valid beats are detected.

##### Stage 1: Detection of questionable beats

The first step of the RCO is the application of AOD on the IBI in both forward and backward direction: those beats that are detected as false positives in both the directions are rejected and those that are detected as true positives in both directions are validated. Each remaining detected beat from backward direction is paired to the nearest detected beat from forward direction. When the distance is less than 250 ms, the pair is set to be checked with the combinatorial optimization.

##### Stage 2: Combinatorial correction

Segments are generated for each sequence of one or more consecutive pairs, by concatenating the last valid beat before the sequence and the first valid beat after the sequence. Then all the combinations of beats are generated by selecting alternatively the beat from the forward or from the backward direction. For each combination we compute the variability index, defined as follows:

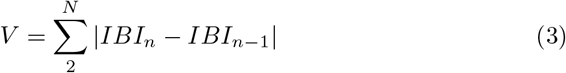

Then the combination with the minimum value of V is selected as the best estimate of beats in the sequence. In the final RCO step, the AOD algorithm is run forward on the corrected IBI to identify possible outliers on IBI series that might have not been corrected.

## 3 Materials and methods

To validate the DBD pipeline we first aimed at testing the performance on a reference dataset. As Physionet^1^ [6] does not provide a dataset of BVP signals with beats annotations, we consider the *Fantasia* dataset [7] which contains the ECG and Blood Pressure (BP) signals (sampling rate: 250 Hz) of twenty subjects (10 young, mean age=27yr and 10 elderly, mean age=74yr) together with beat annotations. Note, however, that BP is a different physiological measure than BVP, though similar. The signals were acquired while the subjects were watching the movie “Fantasia” (Disney, 1940) laying supine and maintaining an inactive state. The heartbeats were first automatically annotated on the ECG and then verified by visual inspection to provide the reference beats series of the Fantasia dataset.

In order to validate the pipeline on signals from WDs, we consider also a dataset appositely created in order to compare the performance between signals from wearable and clinical-grade devices. Physiological data were collected from 18 healthy subjects in two experimental sessions: during resting and during a simulated walking. Both sessions lasted approximately 5 minutes. Each participant gave the informed consent and the experiment was conducted according to the principles of the Declaration of Helsinki [18].

We used a medical-grade device (Biograph) to acquire the reference ECG (2048 Hz, 3 leads configuration) and BVP (2048 Hz). Two WDs were used: the Comftech HeartBand and the Empatica E4. The Comftech HeartBand is a band of elastic tissue with two conductive patches that work as electrodes. The ECG signal is sampled (128 Hz) and amplified by the electronic board which is attached to the band through two snap buttons. The Empatica E4 is a wristband which embeds, among others, a BVP sensor (64 Hz) and a 3-axes accelerometer (32 Hz). Both devices stream the signals through Bluetooth to a smartphone where an appositely developed Android application [2] managed the data flows and sends the compressed data to an online database.

### 3.1 Metrics

We defined two different groups of metrics to quantify performance of the beat detection step and to assess reliability of HRV indicators. The first group compares the results of the beat detection step, the second compares the values of HRV indicators. Four algorithms for beat detection are used: DBD, DBD+RCO, the algorithm for beat detection proposed in [1] with (ABOY+CORR) and without (ABOY) the beat correction procedure.

#### 3.1.1 Beat detection metrics

The results of the beat detection step are validated on the reference beat series first by counting the number of true positives (TPs), false positives (FPs) and false negatives (FNs). To compute the number of TPs, we first try to pair each beat *b_det_* of the detected beat series to a beat *b_ref_* of the reference beats series. The pairing is considered valid when the distance between *b_det_* and *b_ref_* is below 0.5 s. The remaining unpaired detected beats are considered as FPs and the remaining unpaired reference beats are considered as FNs. Then we counted the number of TPs *(n_TP_*), the number of FPs (*n_FP_*) and the number of FNs *((n_FN_*)) to compute the overall *recall ((n_TP_*)/(*n_TP_* + *n_FN_*)) and *precision ((n_TP_)/(n_TP_* + *n_FP_*)).

We also compared the reference IBI series *IBI_ref_* computed from the reference beats series with IBI values resulting from the detected beats *IBI_est_.* We consider the overall root mean square error (rmse) between the interpolated version of *IBI_est_* and *IBI_ref_*. Both the IBI series are interpolated with cubic spline at 4 Hz.

#### 3.1.2 HRV reliability metrics

The second category analyses the reliability of computed HRV indicators. Ten 30-seconds portions of the IBI series are selected and 4 HRV indicators are computed for each portion. Two indicators are chosen from the time-domain category: mean of IBI (RRmean), root mean square of the differences of subsequent IBI (RMSSD), and two from the frequency-domain category: power in the low frequency band [0.04-0.15 Hz] (LF), power in the high frequency band [0.15-0.4 Hz] (HF). The HRV values obtained from the *IBI_ref_* and from the *IBI_est_* are compared using the Bland-Altman (BA) ratio as proposed in [15].

#### 3.1.3 Metrics estimation

Computation of metrics is performed by windowing (length: 30 s, shift: 10 s) along the whole signal, to obtain the distribution of values of each metric for each subject. Beat detection metrics are averaged to obtain the mean value for each subject, while the HRV values are used to compute the BA ratio:

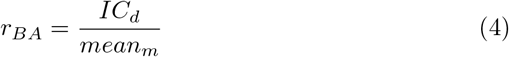

where *IC_d_* is the 95% confidence interval of the difference between the reference and estimated indicator value 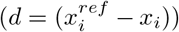, and *mean_m_* is the mean of the average between the reference and estimated indicator value 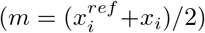. BA ratio is used to evaluate similarity between two sets of measurements. In general, it is used to compare a novel method of measurement with a reference. Two methods are considered as coincident when the BA ratio is below 0.1. A value between 0.1 and 0.2 corresponds to a fair similarity; when the BA ratio is above 0.2, the novel method is considered unreliable.

## 4 Results

### 4.1 Results on Fantasia dataset

For each subject we selected a 1000 s length portion of the BP signal which showed no sensor disconnections. Performance of IBI detection (see Figure 4 appears similar for the DBD and the DBD-RCO pipelines. Performance of ABOY and ABOY-CORR is lower, both in terms of Recall and rmse. Similar results are found also for the Bland-Altman ratios of HRV indicators: DBD and DBD+RCO provide similar results, ABOY and ABOY-CORR are also comparable but value of ratios is higher. However, in general the distribution of the BA values is mostly above the 0.2 value; only for RRmean the ratio is below 0.1 for DBD and DBD-RCO and around 0.2 for ABOY and ABOY-CORR.

**Figure 4:**
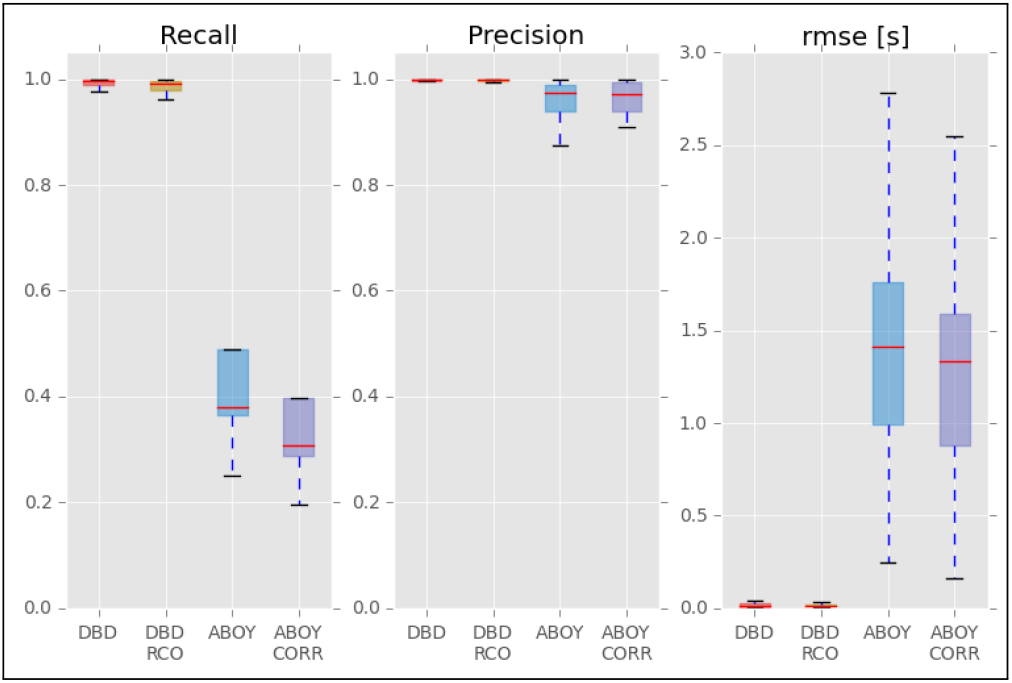
Performances of IBI detection on the Fantasia dataset.

### 4.2 Results on WCS dataset

For each subject we considered both baseline and movement portions to tackle the effect of artifacts. We applied the proposed pipeline to the three cardiac signals acquired during the experiment: (a) BVP signal from the FlexComp, (b) BVP signal from the Empatica E4 and c) the ECG signal from the Comftech HeartBand.

**Figure 5:**
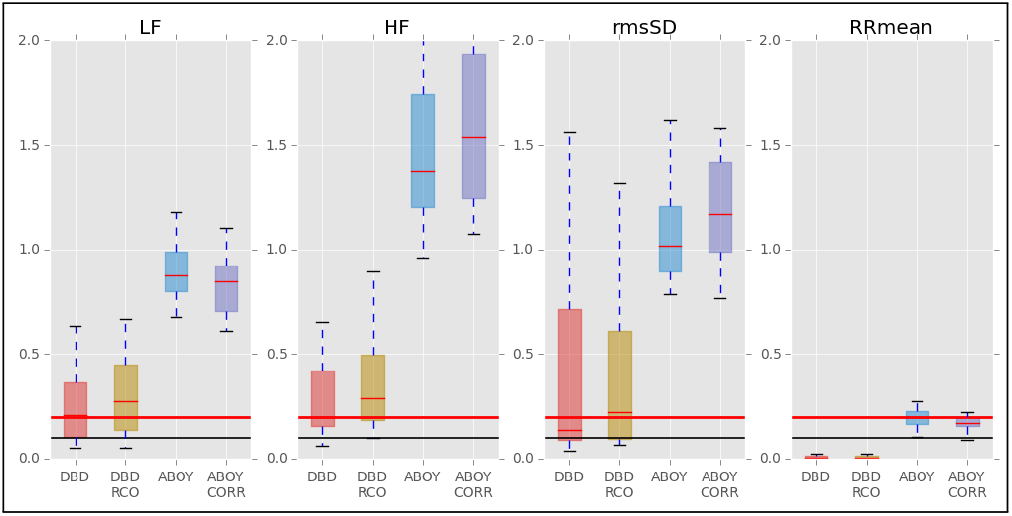
BA ratio of HRV indicators on the Fantasia dataset.

The ECG signal from the FlexComp is used to provide the ground truth: IBI were automatically detected and manually corrected by visual inspection to remove wrong detected peaks and add missing beats.

As with the Fantasia dataset, for the BVP signal (see Figure 6) with the FlexComp and E4 on the baseline the proposed algorithms (DBD and DBD-RCO) perform better than the reference algorithms (ABOY and ABOY-CORR). In general presence of moving artifacts causes an increase in the rmse and decrease of Recall, with more evident effects on the E4.

**Figure 6:**
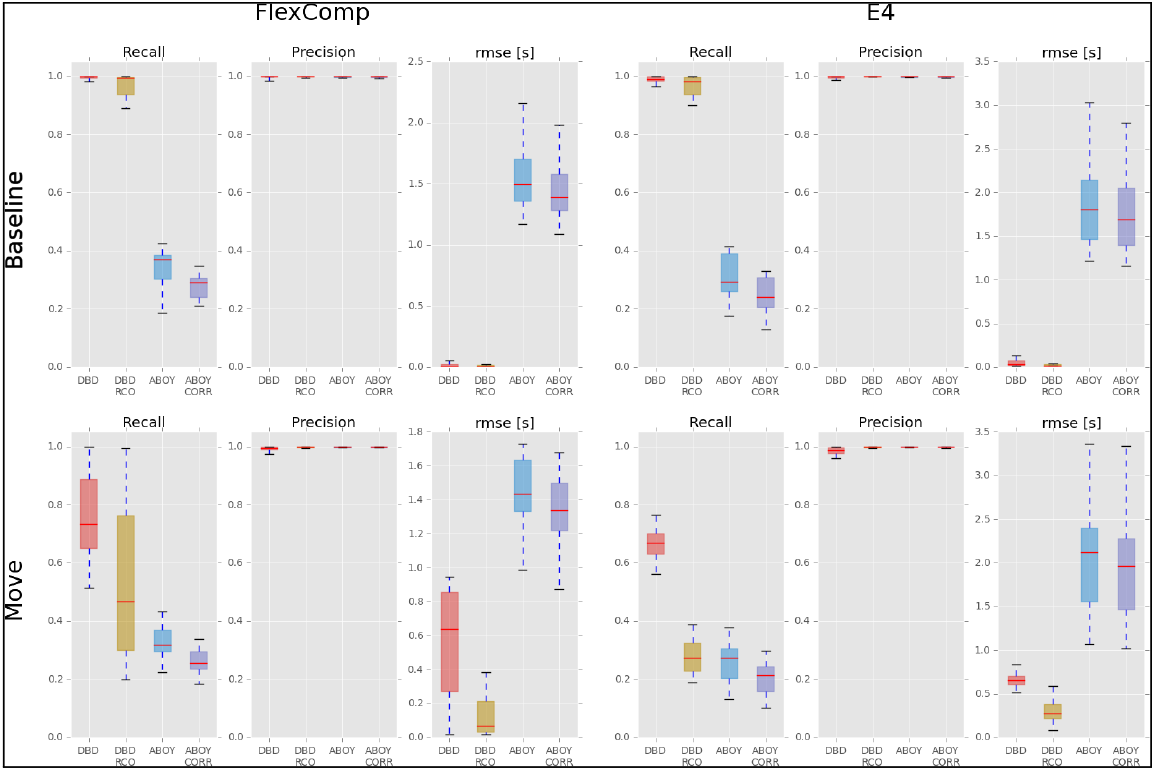
Performances of IBI detection on the BVP from the FlexComp (left) and E4 (right) during baseline (top) and move (bottom) portions.

Reliability indexes of HRV indicators (see Figure 7) extracted from the FlexComp BVP are similar to what found for the Fantasia dataset: better performances of DBD and DBD-RCO algorithms but only RRmean with distribution of BA ratios under 0.2 (around 0 for DBD and DBD-RCO). Results on the BVP from E4 are comparable, although performances of DBD and DBD-RCO are lower.

Body movements cause all the ratios to increase. Again, RRmean from FlexCom appears a reliable indicator, although only the DBD-RCO algorithm presents values below 0.1. For the E4, instead, BA ratio of the RRmean is around 0.2 for all algorithms.

**Figure 7:**
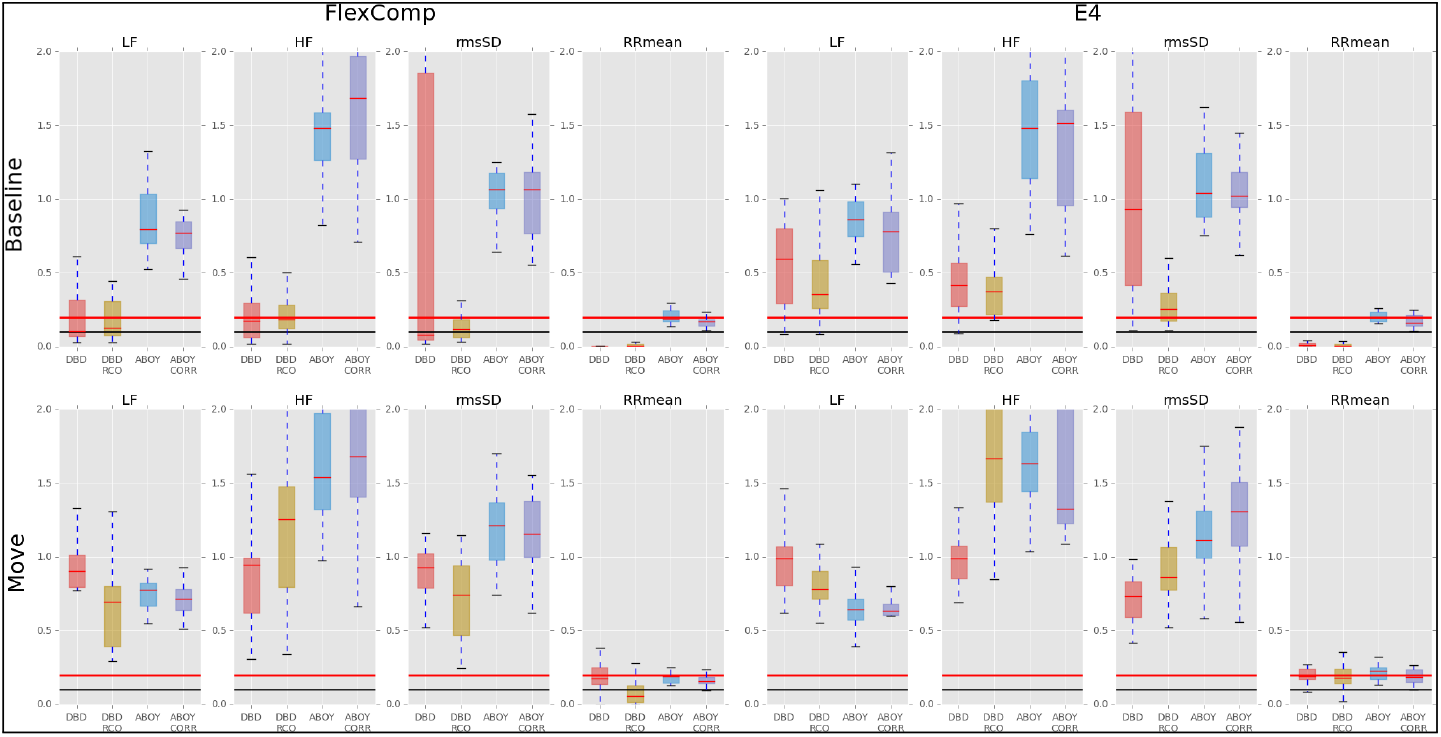
BA ratio of HRV indicators on the BVP from the FlexComp (left) and E4 (right) during baseline (top) and move (bottom) portions. Vertical scale is limited to the interval [0,2] for clarity.

## 5 Discussion

In this paper we proposed the DBD and the RCO a pipeline for detection and processing of cardiac signals from WDs, in particular for beat detection on signals from PPG sensors which represent a common embedded technology to monitor cardiac activity. We compared the proposed algorithm to a reference algorithm [1] in terms of beat detection, differences in the IBI series and HRV indicators, using an existing dataset of clinical signals and a dataset appositely created to assess performances on signals from WDs.

Performance of DBD+RCO improved with respect to the reference algorithm in terms of beat detection and derived IBI values for both the types of devices (clinical and wearable) and both the experimental conditions (baseline and movement). The RCO also improves the results of the beat detection on the ECG signal from the WD Comftech HeartBand.

In absence of body movements the three signals provided similar results in terms of beat detection and IBI series. Regarding HRV indicators the results confirmed that the BVP signals provide unreliable estimates, in particular for frequency domain indicators, as found in other studies [15]. The indicators computed on the Comftech HeartBand ECG signal showed high accuracy in terms of Bland-Altman ratio, however frequency domain indicators presented higher value, but still below the threshold value of 0.2.

During the movements portion the BVP signals from both the FlexComp and the Empatica E4 resulted highly affected by moving artifacts, with slightly better performances for the clinical version (FlexComp). All the HRV indicators, except the RRmean, presented high BA ratio, showing low reliability. Instead, the Comftech HeartBand showed good results in terms of beat detection and IBI series, while, among HRV indicators, only RRmean showed good accordance with the reference.

In summary, the proposed pipeline for processing of cardiac data improves the detection of beats on BVP. In general, cardiac signals acquired with WDs result appropriate to estimate IBI and RRmean, but the errors introduced by the technological constraints of these devices allow no accurate analysis of other HRV indicators. This study highlights also that in presence of body movements an electrocardiography-based device should be used instead of PPG sensors (both clinical and wearable) as it provides a signal which is less sensitive to moving artifacts

## Acknowledgements

This work was supported by a fellowship from TIM, Italy.

## Footnotes

1 http://www.physionet.org

